# Estimation of the reference lead (Pb) concentration levels affecting immune cells in the blood of Black-headed Gulls (*Chroicocephalus ridibundus*, Laridae)

**DOI:** 10.1101/2022.04.27.489755

**Authors:** Nana Ushine, Osamu Kurata, Yoshikazu Tanaka, Shouta M M Nakayama, Mayumi Ishizuka, Takuya Kato, Shin-ichi Hayama

## Abstract

The biological effects of lead (Pb) contamination have been reported in various species. There are no restrictions on the use of Pb products, including bullets, in the areas south of Hokkaido, Japan. Local governments have announced the presence of some Pb in the soil sediments of water bodies. Previous studies have confirmed the relationship between blood Pb level (BLL) and immune cells. This study was performed with the aim of clarifying the effect of Pb contamination on immune cells. In total, 170 Black-headed Gulls (*Chroicocephalus ridibundus*) were captured, including a population in Tokyo Bay between November 2018 and April 2021 and a population in Mikawa Bay between January 2019 and April 2021. Linear regression analysis was performed with the white blood cell count (WBC), proportion of heterophils (Het), proportion of lymphocytes (Lym), ratio of heterophils and lymphocytes (H/L ratio), copy number of CD4 messenger RNA, and copy number of CD8α messenger RNA as the objective variables, and the BLL as the explanatory variable. The group with BLL < 1.0 μg/dL had a significantly lower Het and higher Lym than that with BLL > 3.5 μg/dL (P < 0.05). In addition, the group with BLL < 1.0 μg/dL had a significantly lower H/L ratio than that with BLL > 3.5 μg/dL. CD8α and WBC were higher in the group with the group with BLL range, from 1.0 to 3.5 μg/dL than those in the group with BLL < 1.0 μg/dL. This study suggests that the effect of Pb pollution on the immune cells of Black-headed Gulls is lower than some previous criteria values.

## Introduction

Lead (Pb) is a common environmental pollutant that has physiological effects on many animals [1,2]. In “Risk Communication on Chemical Substances” published by the Ministry of the Environment in Japan, an impact dose is defined as a dose that does not cause any clinical symptoms, a toxic dose causes clinical symptoms and possible death, and a lethal dose has a high probability of death [3]. If a bird ingests a lethal dose, death occurs immediately [4]. Toxic doses cause non-specific symptoms, such as neural symptoms, gastric symptoms, a decline in reproductive success rate, and immune suppression [5,6]. Impact doses also cause non-specific symptoms but these generally have no effect on birds. The symptoms of impact doses are as follows: increased foraging distance from their own population [7], suppression of humoral immunity [8], and increased production of immunoglobulin E (IgE) and cytokines in response to an allergen that causes inflammation [9–12]. Of these impacts, the effect on immunity can lead to mass mortality in the event of highly pathogenic infections in the habitat of Pb-contaminated birds [13]. It is difficult to point out the effects based on impact dose in wildlife, given that no abnormal symptoms are shown [14]. However, impact doses are considered to affect species’ survival as a long-term impact.

Previous research has shown that Black-headed Gulls (*Chroicocephalus ridibundus*, Laridae; winter birds in Honshu, Japan) are contaminated with Pb, and the contamination affects the heterophil and lymphocyte proportions [15]. The purpose of the present study was to determine the minimum Pb level that affects the six cells and molecules in peripheral blood with immunity function, i.e., the total number of white blood cells (WBC; cell/μL), proportion of lymphocytes (Lym; %), proportion of heterophils (Het; %), ratio of heterophils to lymphocytes (H/L ratio), copy number of CD4 messenger RNA (CD4; number of copies/μL), and copy number of CD8α messenger RNA (CD8α; number of copies/μL) in Black-headed Gulls during the winter season. This study contributes to the risk assessment of wild bird pathogens by clarifying the effects of Pb pollution on wild birds and their immune function on cells and molecules.

## Materials and methods

### Ethics statement

This field study was approved by the Ministry of the Environment, Chiba Prefecture (approval number: 739-1568, 1667, 900), and Aichi Prefecture (approval number: 588-5, 1013-3). This study was assessed the degree of distress suffered by gulls, and all work under investigation was reviewed and approved by the Laboratory Animal Ethics Committee at Nippon Veterinary and Life Science University (approval number: 30S-47, 2020K-78).

### Sample animals

Black-headed Gulls were captured by hand, noose trap, or whoosh net from November 2018 to April 2021 at Tokyo Bay (N35°, E139°) and from January 2019 to April 2021 at Mikawa Bay (N34°, E137°). Peripheral blood (less than 0.5% of body mass) was collected using a heparin-coated syringe (Nipro, Osaka, Japan) according to the method of Gaunt et al. [16]. After hemostasis was confirmed, a color ring and a metal ring for bird banding were attached to the left and right tarsus of the gulls in the Tokyo Bay population, respectively, and only a metal ring was attached to the right tarsus of the Mikawa Bay gulls.

### Identification of sex and age

Age was assessed using the plumage and birds were classified into two groups: yearlings (which had first-year plumage) and adults (with no first-year plumage) [17]. Sex was identified by molecular methods using a chromodomain helicase DNA-binding protein gene on the sex chromosomes as the target gene [18]. The protocols for the polymerase chain reaction (PCR) and primer design by Ushine et al. [19] were followed.

### Evaluation of immunity

The following six indices of peripheral blood were used to evaluate immunity [15]: WBC was counted (cell number per deciliter), Lym and Het were calculated as the ratio of Lym and Het per 100 leukocytes using a smear stained with light Giemsa, and the H/L ratio was calculated as a ratio of these results. The number of copies of CD4 and CD8α cells in peripheral blood was calculated using real-time PCR using primers specific to Black-headed Gulls.

### Measurement of Pb level

Clots in the peripheral blood were autoclaved and transported to Hokkaido University (Kita 8, Nishi 5, Kita-ku, Sapporo, Hokkaido, Japan). The blood Pb level (BLL) was measured using the method of Ushine et al. [13] and converted to the amount of Pb per deciliter of a clot.

### Statistical analysis

Pb toxicity causes different symptoms depending on the concentration of the pollutant in each species [20]. In general, the reference concentration of Pb contamination is 20 μg/dL in birds [21]. The reference values of BLL have been advocated in some avian groups that have already been reported to be affected by severe poisoning or pollution [22]. For example, the concentration of Pb in the order Anseriformes is reported to be 20 μg/dL [23] and that in the genus *Haliaeetus* is reported to be 40 μg/dL [24].

However, establishing the threshold levels as reference values for heavy metal pollution requires exposure experiments using animals [25]. It is difficult to set a threshold for Black-headed Gulls because no captive exposure experiments have been performed for this species. Thus, in this study, the BLL was classified into quartile following the methodology of Krishnan et al. [26]. The BLL samples were grouped as follows: < 1.0 μg/dL, 1.0 to 2.0 μg/dL, 2.0 to 3.5 μg/dL, and > 3.5 μg/dL.

First, differences in Pb levels with respect to age, sex, and population were confirmed using the Wilcoxon rank-sum test. If these indices had any significant differences, the gull populations were classified according to these indices. Next, the Shapiro–Wilk test was used to confirm that the objective variables WBC, Het, Lym, H/L ratio, CD4, and CD8α were normally distributed. To obtain normal distributions, CD4 and CD8α were logarithmically converted before analysis. Finally, linear regression analysis was performed with BLL as the explanatory variable. The group with the lowest BLL (< 1.0 μg/dL) was used as a reference. Statistical analyses were performed using R software (ver. 3.5.0) and Stata (ver. 14.0). The significance level was 0.05 for all statistical analyses.

## Results

A total of 170 gulls were captured and blood samples were analyzed (Table 1). Differences in Pb contamination by age, sex, and population were not significant, and block structures for these variables were not included. The result value of six indices in peripheral blood and BLL is shown in Table 2. Linear regression analysis revealed significant trends in five indices (Table 3). First, the group with BLL < 1.0 μg/dL had lower Het than that with BLL > 3.5 μg/dL (P < 0.05). Second, the group with BLL < 1.0 μg/dL had higher Lym than that with BLL > 3.5 μg/dL (P < 0.05; Fig. 1). The H/L ratio was significantly higher in the group with BLL > 3.5 μg/dL than in that with BLL < 1.0 μg/dL (Fig. 2). WBC (Fig. 3) and CD8α (Fig. 4) were significantly higher in the groups with BLL 1.0 to 2.0 and 2.0 to 3.5 μg/dL than in that with BLL < 1.0 μg/dL (P < 0.05).

**Table 1.** Results of blood test items in 170 Black-headed Gulls (*Chroicocephalus ridibundus*) comprising a Tokyo Bay group captured from November 2018 to April 2021 and a Mikawa Bay group captured from January 2019 to April 2021.

| Items | Pb groups (sample size) | Coef. | P value | 95 % CI |
| --- | --- | --- | --- | --- |
| WBC | < 1.0 µg/dL (43) | Reference |  |  |
|  | 1.0 to 2.0 µg/dL (52) | 1586.05 | 0.01 | 332.65 – 2839.45 |
|  | 2.0 to 3.5 µg/dL (34) | 2456.64 | 0.001 | 1061.11 – 3852.15 |
|  | > 3.5 µg/dL (41) | 1264.10 | 0.06 | -63.23 – 2591.42 |
| Het | < 1.0 µg/dL (43) | Reference |  |  |
|  | 1.0 to 2.0 µg/dL (52) | 1.49 | 0.38 | -1.87 – 4.85 |
|  | 2.0 to 3.5 µg/dL (34) | 2.81 | 0.14 | -0.94 – 6.55 |
|  | > 3.5 µg/dL (41) | 8.68 | 0.0001> | 5.12 – 12.25 |
| Lym | < 1.0 µg/dL (43) | Reference |  |  |
|  | 1.0 to 2.0 µg/dL (52) | -0.76 | 0.65 | -4.11 – 2.59 |
|  | 2.0 to 3.5 µg/dL (34) | -3.36 | 0.08 | -7.09 – 0.37 |
|  | > 3.5 µg/dL (41) | -8.34 | 0.0001> | -11.89 – -4.79 |
| H/L ratio | < 1.0 µg/dL (43) | Reference |  |  |
|  | 1.0 to 2.0 µg/dL (52) | 0.04 | 0.43 | -0.06 – 0.15 |
|  | 2.0 to 3.5 µg/dL (34) | 0.11 | 0.07 | -0.01 – 0.22 |
|  | > 3.5 µg/dL (41) | 0.29 | 0.0001> | 0.18 – 0.40 |
| CD4 | < 1.0 µg/dL (43) | Reference |  |  |
|  | 1.0 to 2.0 µg/dL (52) | 0.28 | 0.07 | -0.02 – 0.59 |
|  | 2.0 to 3.5 µg/dL (34) | -0.01 | 0.94 | -0.35 – 0.32 |
|  | > 3.5 µg/dL (41) | 0.25 | 0.12 | -0.07 – 0.57 |
| CD8α | < 1.0 µg/dL (43) | Reference |  |  |
|  | 1.0 to 2.0 µg/dL (52) | 1.36 | 0.001 | 0.55 - 2.17 |
|  | 2.0 to 3.5 µg/dL (34) | 3.14 | 0.0001> | 2.24 - 4.05 |
|  | > 3.5 µg/dL (41) | 0.07 | 0.87 | -0.78 - 0.93 |

**Table 2.**
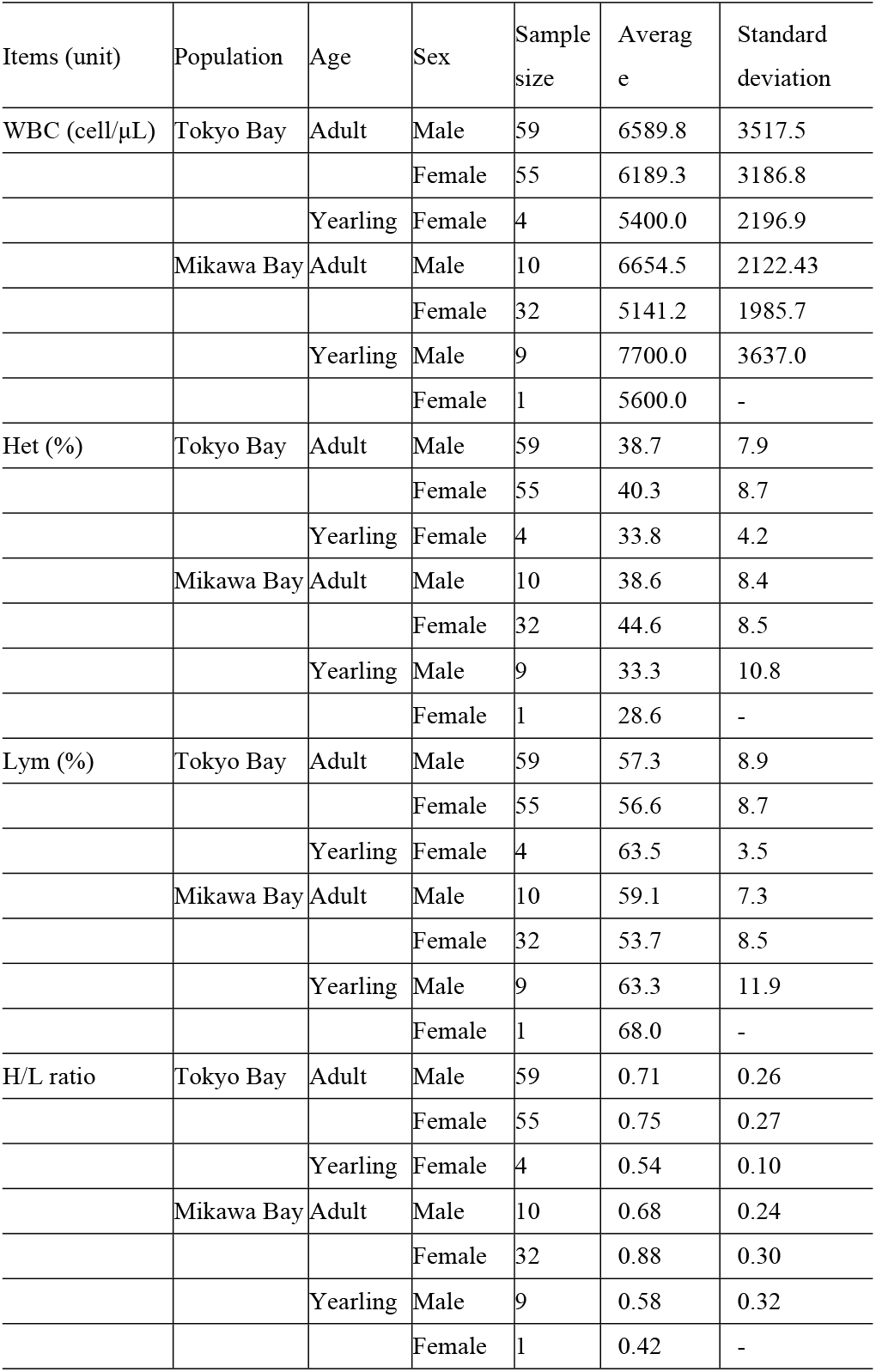
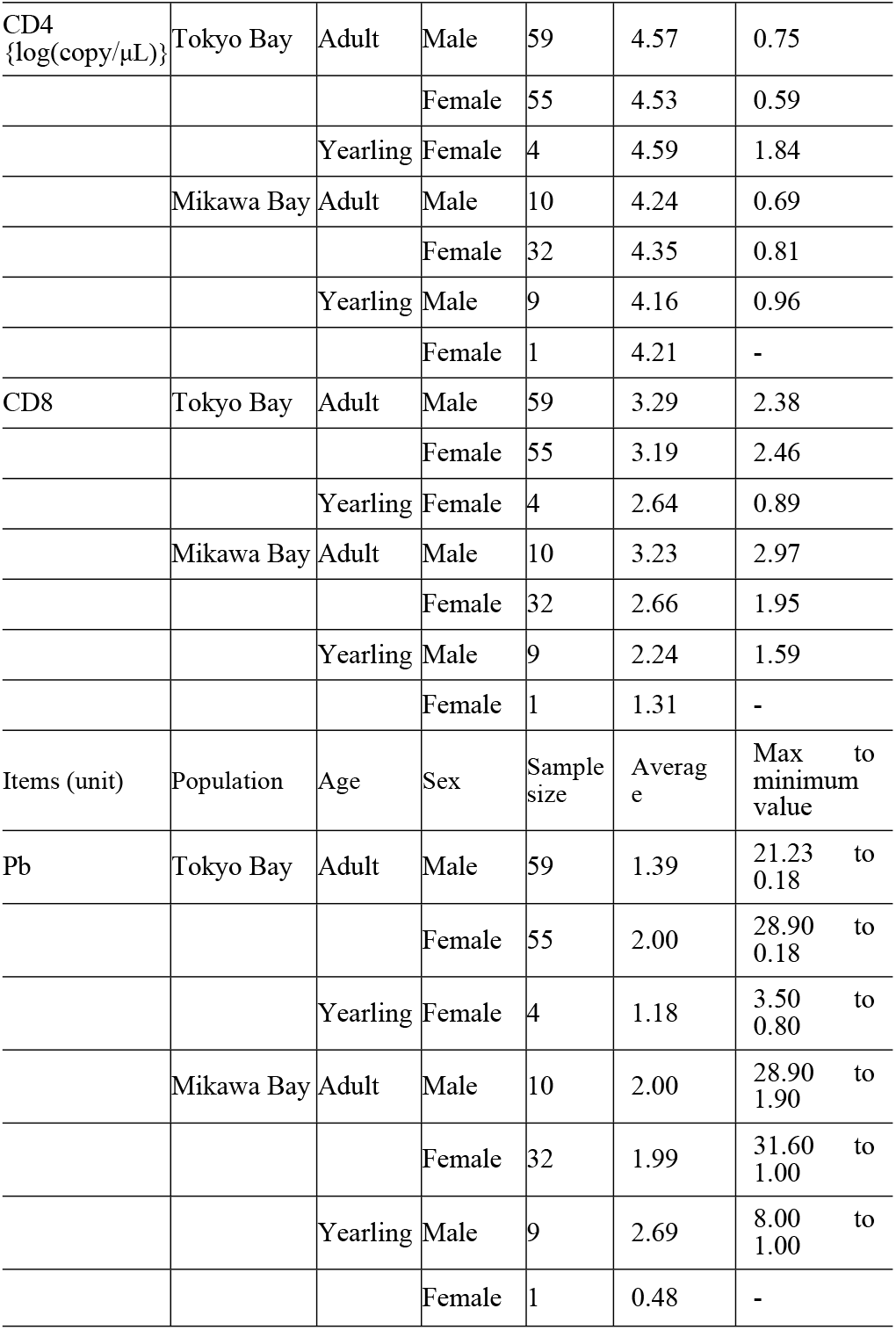
Results of blood test variables when blood Pb level is classified into quartiles. in 170 Black-headed Gulls (*Chroicocephalus ridibundus*) comprising a Tokyo Bay group captured from November 2018 to April 2021 and a Mikawa Bay group captured from January 2019 to April 2021.

**Table 3.** Linear regression analysis of immunity parameters classified into quartiles in blood Pb level (BLL) in 170 black-headed gulls (*Chroicocephalus ridibundus*) comprising a Tokyo Bay group captured from November 2018 to April 2021 and a Mikawa Bay group captured from January 2019 to April 2021. The group with BLL < 1.0 μg/dL was used as a reference group. Abbreviations: Coef. = coefficient of determination, 95% CI = 95% confidence interval, WBC = total number of white blood cells, Het = proportion of heterophils, Lym = proportion of lymphocytes, H/L ratio = ratio of heterophils to lymphocytes, CD4 = copy number of CD4 messenger RNA, CD8α = copy number of CD8α messenger RNA.

| Quartile of BLL ($\mu\text{g/dL}$) | <1.0 | 1.0 to 2.0 | 2.0 to 3.5 | > 3.5 |
| --- | --- | --- | --- | --- |
| Sample size | 43 | 52 | 34 | 41 |
| WBC |  |  |  |  |
| Average | 4814 | 6400 | 7271 | 6078 |
| $\pm 1$ SD | 2947 | 3083 | 3063 | 3224 |
| Het |  |  |  |  |
| Average | 37.2 | 38.7 | 40.0 | 45.9 |
| $\pm 1$ SD | 7.6 | 8.2 | 9.1 | 8.4 |
| Lym |  |  |  |  |
| Average | 59.6 | 58.8 | 56.2 | 51.2 |
| $\pm 1$ SD | 7.6 | 8.1 | 9.3 | 8.1 |
| H/L ratio |  |  |  |  |
| Average | 0.65 | 0.69 | 0.76 | 0.94 |
| $\pm 1$ SD | 0.20 | 0.24 | 0.28 | 0.30 |
| CD4 |  |  |  |  |
| Average | 4.33 | 4.61 | 4.32 | 4.58 |
| $\pm 1$ SD | 0.62 | 0.84 | 0.76 | 0.72 |
| CD8α | | | | |
| Average | 2.00 | 3.36 | 5.14 | 2.08 |
| $\pm 1$ SD | 0.63 | 2.22 | 3.37 | 0.64 |

**Figure 1.**
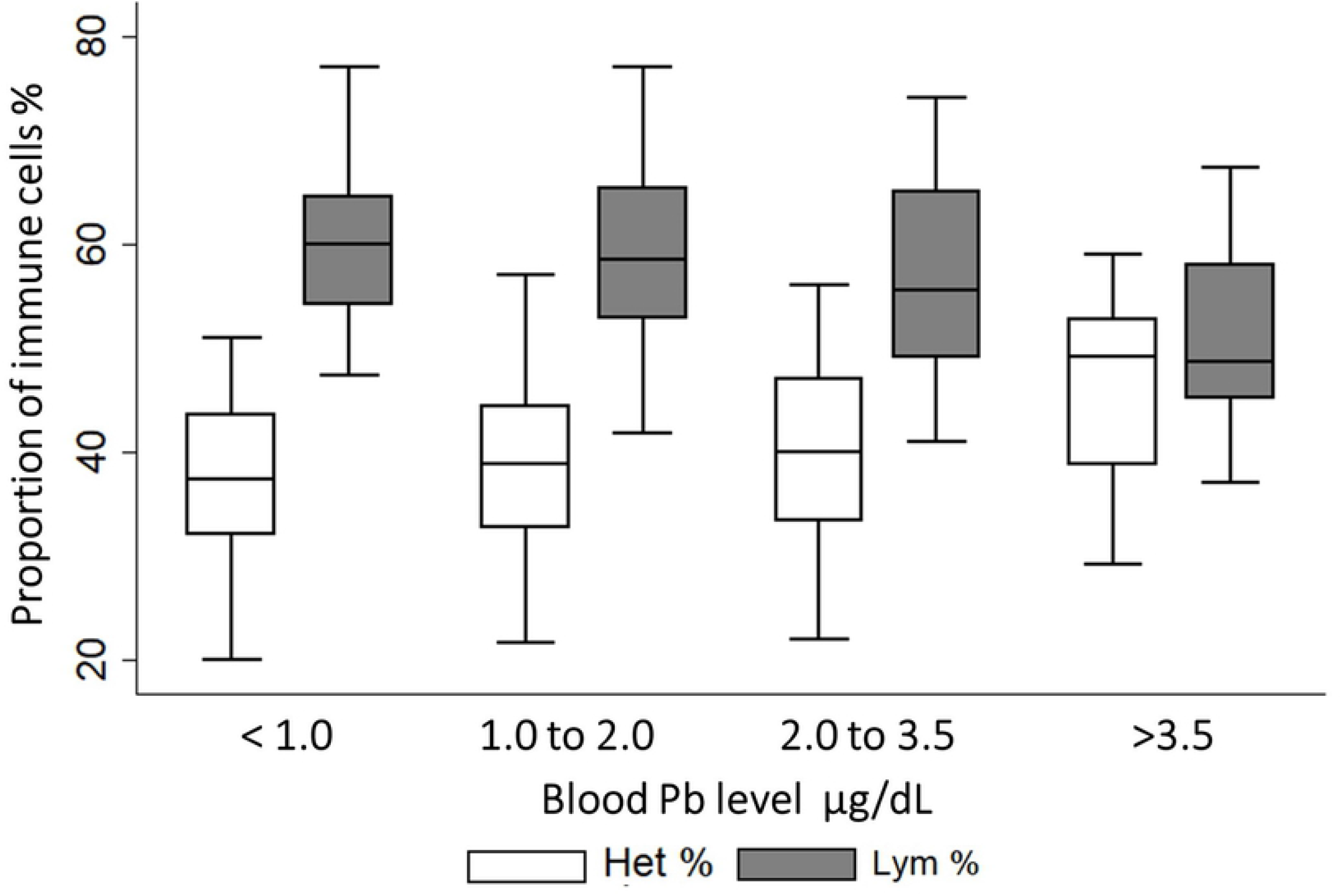
Linear regression analysis of the heterophil to lymphocyte ratio for quartiles in blood Pb level (BLL) in 170 Black-headed Gulls (*Chroicocephalus ridibundus*) comprising a Tokyo Bay group captured from November 2018 to April 2021 and a Mikawa Bay group captured from January 2019 to April 2021. The group with BLL < 1.0 μg/dL was used as a reference group. The center bar shows the median (%), and the upper and lower bars show the maximum and minimum values of the boxplot, respectively. Dots indicate outliers. The proportion of lymphocytes (Lym) in the group with BLL < 1.0 μg/dL was significantly higher than that in the group with BLL > 3.5 μg/dL. The proportion of heterophils (Het) was significantly lower in gulls with BLL < 1.0 μg/dL (P < 0.05).

**Figure 2.**
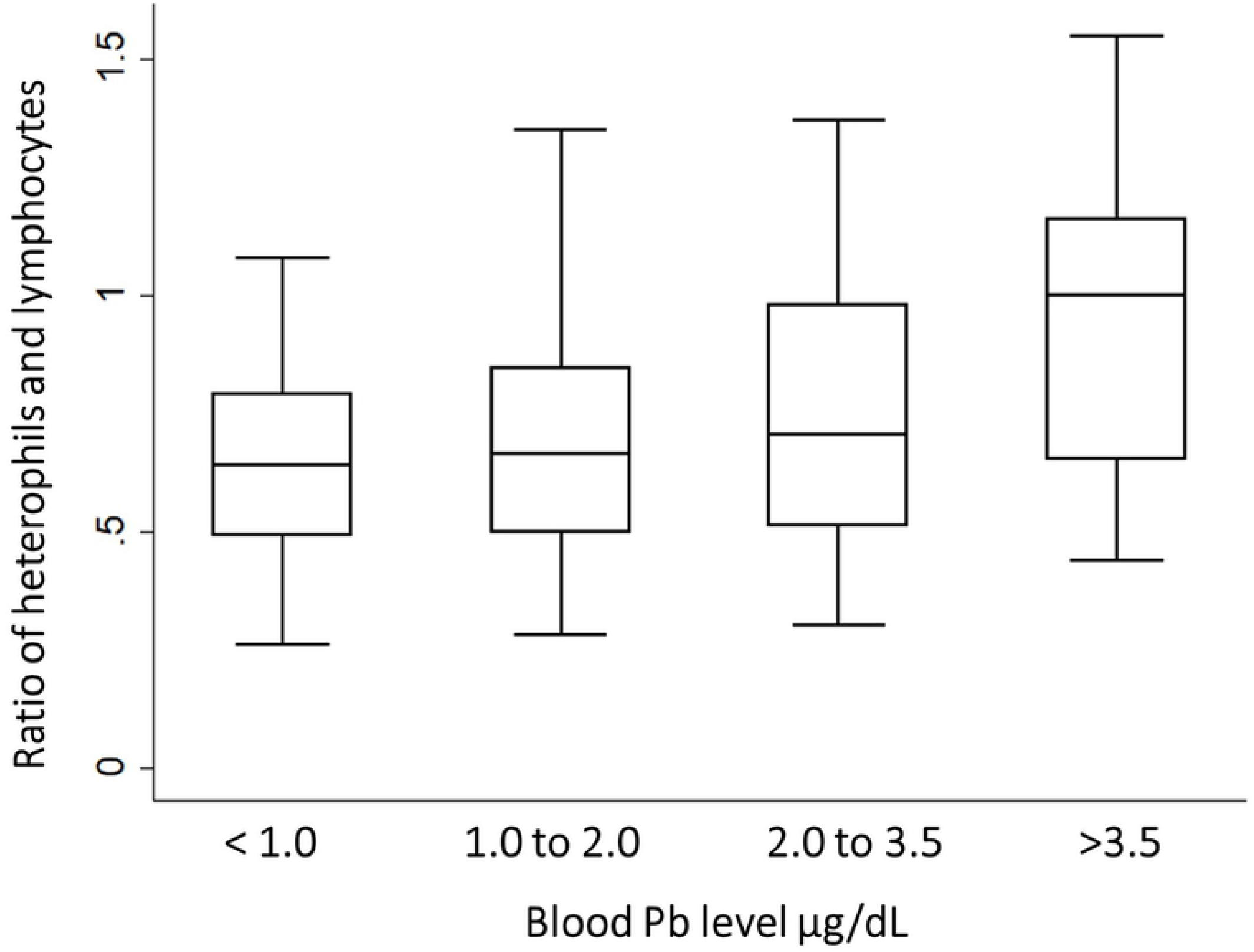
Linear regression analysis of the heterophil to lymphocyte (H/L) ratio for increments in blood Pb level (BLL) of quartile in 170 Black-headed Gulls (*Chroicocephalus ridibundus*), comprising a Tokyo Bay group captured from November 2018 to April 2021 and a Mikawa Bay group captured from January 2019 to April 2021. The group with BLL < 1.0 μg/dL was used as a reference group. The center bar shows the median (%), and the upper and lower bars show the maximum and minimum values of the boxplot, respectively. Dots indicate outliers. The H/L ratio for the groups with BLL < 1.0 μg/dL was lower than that for the group with BLL > 3.5 μg/dL (P < 0.05).

**Figure 3.**
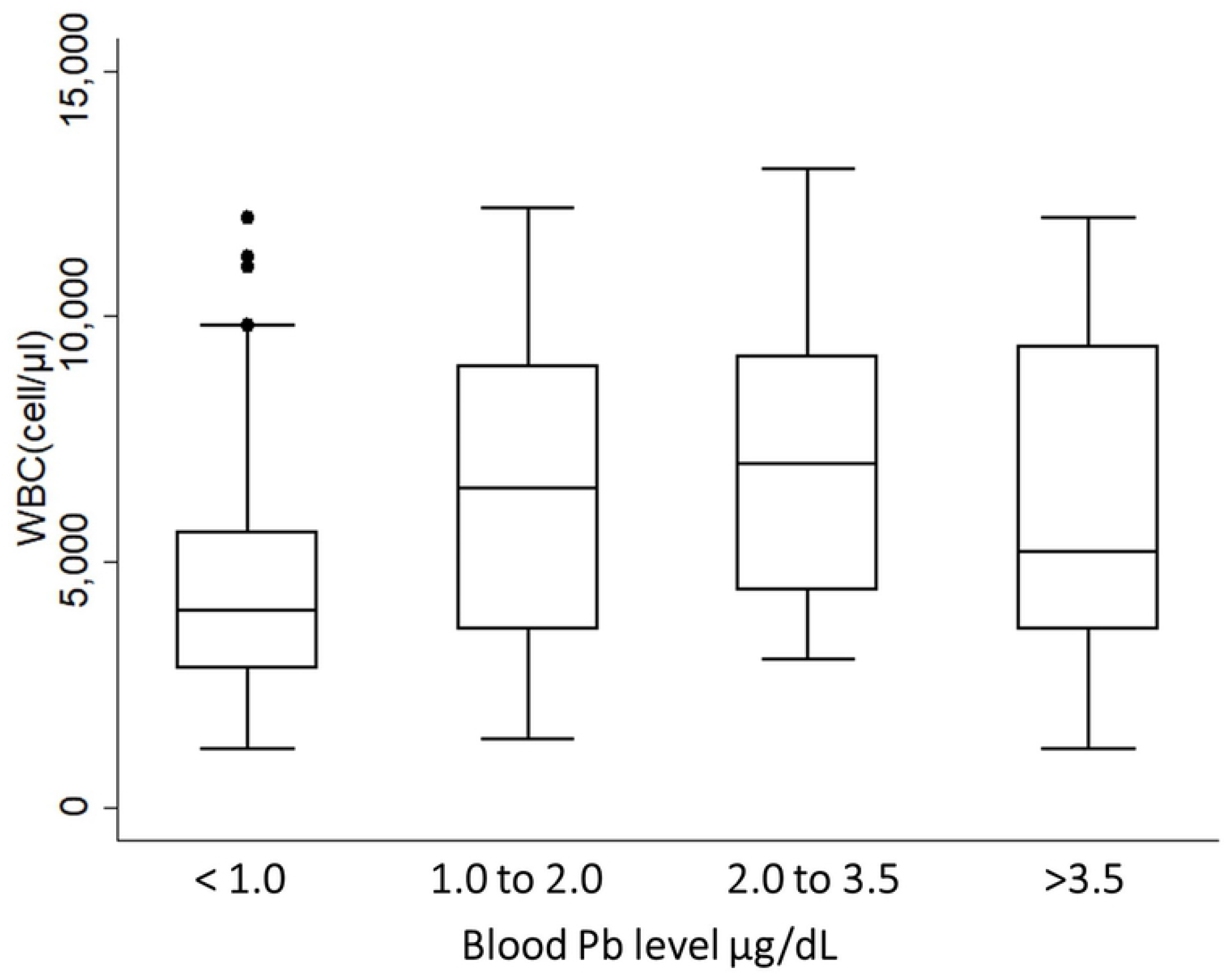
Linear regression analysis of the number of white blood cell (WBC) for increments in blood Pb level (BLL) of quartile in 170 Black-headed Gulls (*Chroicocephalus ridibundus*), comprising a Tokyo Bay group captured from November 2018 to April 2021 and a Mikawa Bay group captured from January 2019 to April 2021. The group with BLL < 1.0 μg/dL was used as a reference group. The center bar shows the median (%), and the upper and lower bars show the maximum and minimum values of the boxplot, respectively. Dots indicate outliers. The WBC for the group with BLL < 1.0 μg/dL was lower than that for the group with BLL 1.0 to 3.5 μg/dL (P < 0.05).

**Figure 4.**
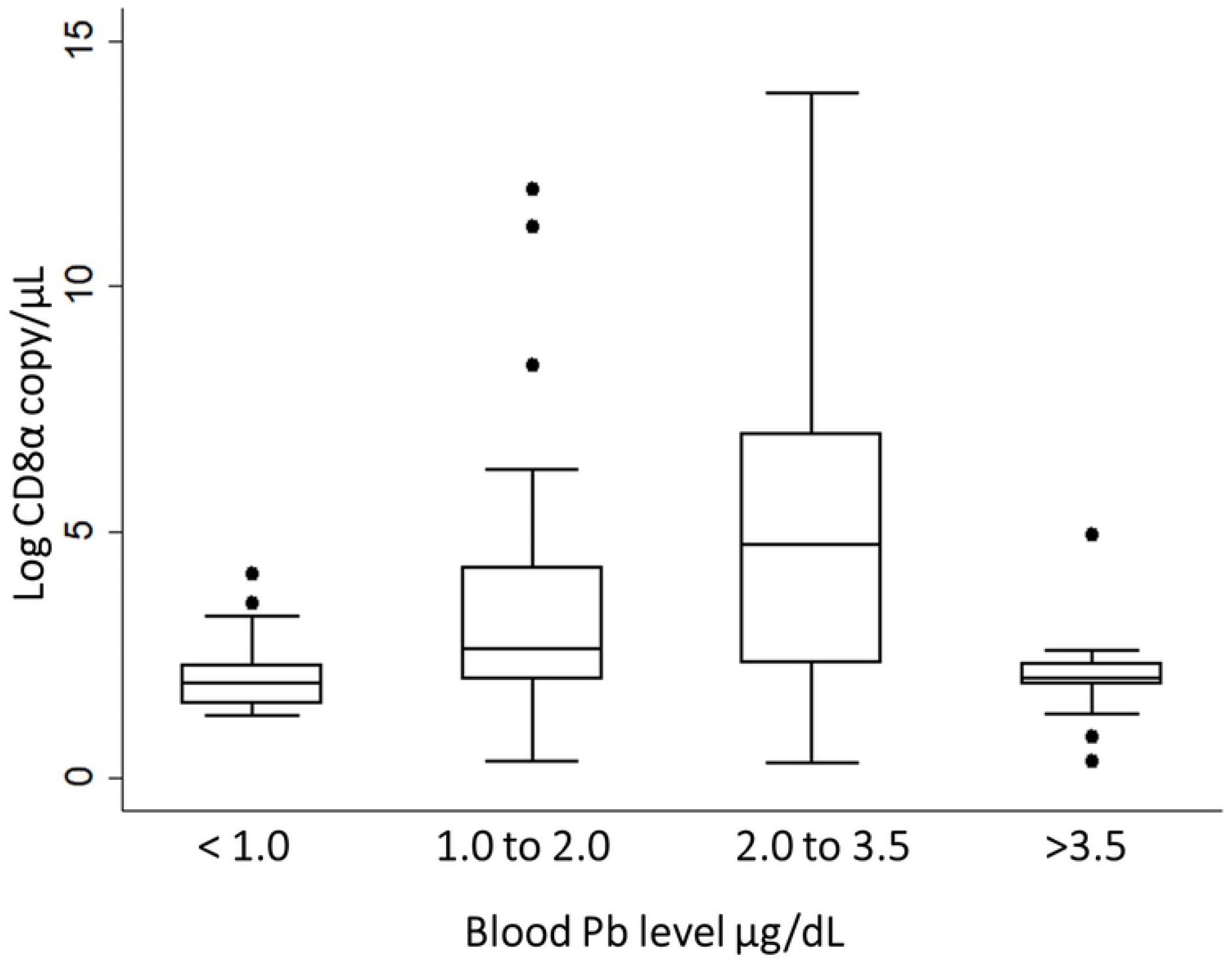
Linear regression analysis of the copy number of CD8α messenger RNA (CD8α) for increments in blood Pb level (BLL) of quartile in 170 Black-headed Gulls (*Chroicocephalus ridibundus*), comprising a Tokyo Bay group captured from November 2018 to April 2021 and a Mikawa Bay group captured from January 2019 to April 2021. The group with BLL < 1.0 μg/dL was used as a reference group. The center bar shows the median (%), and the upper and lower bars show the maximum and minimum values of the boxplot, respectively. Dots indicate outliers. The CD8α for the group with BLL < 1.0 μg/dL was lower than that for the group with BLL 1.0 to 3.5 μg/dL (P < 0.05).

## Discussion

In this study, the proportion of Het increased for BLL > 3.5 μg/dL, whereas Lym decreased at the same concentration. An increase in Het affects the suppression of the immune response caused by the inflammatory response [27] and a decrease in Lym has a negative effect on the regulation of immune function [28]. Therefore, when the BLL > 3.5 μg/dL, the Het and Lym levels changed substantially, leading to possible negative effects on immune function. Similar to these results, the H/L ratio showed a significant increase > 3.5 μg/dL. The H/L ratio is maintained at a certain value for each avian species [29]. Deviations in the H/L ratio indicate abnormal conditions, such as stress and starvation [30]. The current study suggests that the proportions of Het and Lym were affected at BLL > 3.5 μg/dL as a result of increases and decreases in the levels of the two types of cells that configure this ratio.

WBC and CD8α were significantly higher in the group with BLL 1.0 to 3.5 μg/dL than in that with BLL <1.0 μg/dL, unlike the results of Het and Lym. The biological effect of Pb pollution is dependent on the Pb level and dose per unit time [31]. The herring gull (*Larus argentatus*) shows a 16 % reduction in equilibrium capacity after exposure to 100 mg/kg Pb compared with that to 50 mg/kg Pb [32]. Fritsh et al. [33] reported that the reproductive success rate of the Black-bird (*Turdus merula*) polluted with under 10 μg Pb/g dry weight is reduced. In some avian species, low Pb pollution has been reported to reduce bodily strength [5,34]. The changes in WBC and CD8α in response to Pb have been shown for the first time in the current study and is considered to depending on the level of the pollutant.

In the current study, only CD4 was not significantly affected by BLL. CD4 positive cells are the most affected by Pb pollution in immune cells [35]. Cao et al. [36] found that the number of CD4 memory cell are increased by Pb contamination, whereas those of naive CD4 cells are decreased. This report effect is considered to be one of the reasons why this study did not show a significant relationship between BLL and CD4; another potential cause is that the Pb level in this study was less than the level that affects CD4. Pb contamination in workers is reported to contribute to the activation of Th2 cells involved in the proliferation of Th1 cells, which are CD4+ T cells [35]. In contrast, children with a high BLL have a significantly lower proportion of CD4+ cells [36]. In the current study, the Black-headed Gulls did not develop any toxicity symptoms, implying that they were chronically polluted with Pb at an impact dose. In addition, the effect of Pb differs depending on the type of cells expressing CD4 [37]; it is, therefore, possible that no significant change was observed in the CD4 level. In a 30,000 μg/dL Pb exposure experiment in rats, the CD4 level was significantly decreased, whereas that of CD8 was not changed [38]. Considering the results of these studies, it was concluded that there was no clear relationship between CD4 and BLL in gulls.

Although the mechanism of Pb contamination on avian immune suppression has not been elucidated, some case studies report immunosuppression after Pb contamination. For example, in helper T cells, which express CD4 on the surface, immune activity is significantly suppressed at a blood value ≥ 10 μg/dL in juvenile Mallard (*Anas platyrhynchos*) [39]. In a Pb exposure experiment in Japanese quail (*Coturnix japonica*), the expression level of Toll-like receptor-3, which has an antiviral effect, was significantly decreased in the group exposed with BLL of 5,000 μg/dL than in that with BLL of 500 μg/dL [40]. For some bird groups, reference values for pollution have been set based on the concentration at which symptoms of poisoning are observed, such as in Anseriformes [23] and *Haliaeetus* species [24]. The biological effects of low Pb level pollution have been investigated in detail in children, thus in 2012, the CDC set a standard value of 5 μg for Pb contamination [41]. However in 2021, the CDC was revised to 3.5 μg due to the impact it had on the intellectual development of infants [42]. Although the effect of BLL is mediated by immune cells in animal species, the BLL that affects immunity in Black-headed Gulls is lower than that for other species. For example, in Mallards, chronic contamination with Pb has been shown to significantly increase the infection rate and species number of intestinal parasites [43]. In addition, a positive relationship was reported between the parasitism rate of *Plasmodium relictum* and Pb level in feathers in the house sparrow (*Passer domesticus*) [44]. None of the birds in these reports show any sign of poisoning, suggesting that an impact dose of Pb increases the risk of infection. Therefore, there is a concern that the BLL revealed in this research may increase the risk of infection by pathogens.

## Conclusion

This study revealed that even the migratory Black-headed Gulls that inhabit Japan only for the winter season are contaminated with Pb, and that BLL > 3.5 μg/dL affects the proportions of heterophils and lymphocytes, that contribute to immune function. Moreover, it was confirmed that the number of WBC and copy number of CD8α messenger RNA, which are related to immune function, increased significantly at BLL 1.0 to 3.5 μg/dL. The results of this study suggest that significant changes in the proportion of some blood cells and molecules with immune function occur owing to Pb contamination. This occurs at a dose that is not toxic but considered an impact dose. If the gull continues to inhabit polluted areas, it is more likely to be infected by pathogens or contaminated with Pb in the wintering areas.

## Acknowledgments

We thank Mr. Yoshihiro Kurahashi, Mr. Tatsuo Sato, and all staff of the Gyotoku Bird Observatory Society NPO. We are grateful to Ms. S. Hirota, Ms. H. Sasaki, Ms. H. Otsubo, Dr. S. Moriguchi, and Dr. N. Sugiura from the Department of Wildlife medicine of the Nippon Veterinary and Life Science University of Tokyo prefecture for the great cooperation in this survey. We also thank to Dr. M. Onuma from the Biodiversity division of the National Institute for Environmental Studies, Japan, Dr. T. Yamamoto from the Department of Veterinary Nursing and Technology of the Nippon Veterinary and Life Science University for many technical advises. Finally, we are grateful to Ms. N. Hirano amd Mr. T. Ichise for the critical help with lead measurement. This work was supported by the Sasakawa Scientific Research Grant from The Japan Science Society and the Environment Research and Technology Development Fund (SII-1) of the Environmental Restoration and Conservation Agency of Japan. This research was also supported by JST/JICA, SATREPS (Science and Technology Research Partnership for Sustainable Development; No. JPMJSA1501) and Program for supporting introduction of the new sharing system (JPMXS0420100619). This work was supported by the Grants-in-Aid for Scientific Research from the Ministry of Education, Culture, Sports, Science and Technology of Japan awarded to S.M.M. Nakayama (20K20633).

## Notes

### Competing Interest Statement

The authors have declared no competing interest.

## References

1. Cromie R, Newth J, Reeves J, O’Brien M, Beckman K, Brown M. The sociological and political aspects of reducing lead poisoning from ammunition in the UK: why the transition to non-toxic ammunition is so difficult. Oxford Lead Symposium. 2014. December 10 [Cited 2020 June 1] Available from: http://www.oxfordleadsymposium.info/wp-content/uploads/OLS_proceedings/papers/OLS_proceedings_cromie_newth_reeves_obrien_beckman_brown.pdf

2. Masindi V, Muedi KL. Environmental contamination by heavy metals. In: Saleh HE-DM, Aglan RF, editors. Heavy Metals. London: Intech Open; 2018. pp 115–133.

3. Ministry of the Environment: Chemicals Advisor Certification Examination Text. (In Japanese) 2008 April 1 [Cited 2020 December 11] Available from: http://www.env.go.jp/chemi/communication/taiwa/text/3s.pdf.

4. Pain DJ, Fisher I, Thomas VG. A global update of lead poisoning in terrestrial birds from ammunition sources. In R. T. Watson, M. Fuller, M. Pokras, and W. G. Hunt, editors. Ingestion of Lead from Spent Ammunition: Implications for Wildlife and Humans. The Peregrine Fund: Boise. 2009.

5. Kendall RJ, Lacher Jr. TE, Bunck C, Daniel B, Driver C, Grue CE, et al. An ecological risk assessment of lead shot exposure in non-waterfowl avian species: upland game birds and raptors. Environ Toxicol Chem. 1996; 15: 4–20. https://doi.org/10.1002/etc.5620150103

6. Williams RJ, Holladay SD, Williams SM, Gogal Jr. RM. Environmental Lead and Wild Birds: A Review. Rev Environ Contam Toxicol. 2018; 245: 157–180. https://doi.org/10.1007/398_2017_9

7. Pain DJ, Mateo R, Green RE. Effects of lead from ammunition on birds and other wildlife: a review and update. Ambio. 2019; 48: 935–953. https://doi.org/10.1007/s13280-019-01159-0

8. Snoeijs T, Dauwe T, Pinxten R, Vandesande F, Eens M. Heavy metal exposure affects the humoral immune response in a free-living small songbird, the great tit (*Parus major*). Arch Environ Contam Toxicol. 2014; 46: 399–404. https://doi.org/10.1007/s00244-003-2195-6

9. Heo Y, Lee WT, Lawrence DA. In vivo the environmental pollutants lead and mercury induce oligoclonal T cell responses skewed toward Type-2 reactivities. Cell Immunol. 1997; 179: 185–195. https://doi.org/10.1006/cimm.1997.1160

10. Miller T, Golemboski K, Ha R, Bunn T, Sanders F, Dietert R. Developmental exposure to lead causes persistent immunotoxicity in Fischer 344 rats. Toxicol Sci. 1998; 42: 129–135. http://dx.doi.org/10.1006/toxs.1998.2424

11. Tepper RI, Levinson DA, Stanger BZ, Campos-Torres J, Abbas AK, Leder P. IL-4 induces allergic-like inflammatory disease and alters T cell development in transgenic mice. Cell. 1990; 62: 457–467. https://doi.org/10.1016/0092-8674(90)90011-3

12. Wood N, Bourque K, Donaldson DD, Collins M, Vercelli D, Goldman SJ, Kasaian MT. IL-21 effects on human IgE production in response to IL-4 or IL-13. Cell Immunol. 2004; 231: 133–145. https://doi.org/10.1016/j.cellimm.2005.01.001

13. Ushine, N, Nakayama SMM, Ishizuka M, Sato T, Kurahashi Y, Wakayama E, Sugiura N, Hayama S. Relationship between blood test values and blood lead (Pb) levels in Black-headed gull (*Chroicocephalus ridibundus*: Laridae). J Vet Med Sci. 2020a; 82: 1124–1129. https://doi.org/10.1292/jvms.20-0246

14. Johnson CK, Vodovoz T, Boyce WM, Mazet JA. Lead exposure in California condors and sentinel species in California. Report prepared for the California Department of Fish and Game. 2007 February 14 [Cited 2021 April 4] Available from: https://citeseerx.ist.psu.edu/viewdoc/download?doi=10.1.1.523.6628&rep=rep1&type=pdf#:~:text=Lead%20toxicity%20is%20believed%20to,(Sorenson%20and%20Burnett%202007)

15. Ushine N, Kurata O, Tanaka Y, Sato T, Kurahashi Y, Hayama S. The effects of migration on the immunity of Black-headed gulls (*Chroicocephalus ridibundus*: Laridae). J Vet Med Sci. 2020b; 82: 1619–1626. https://doi.org/10.1292/jvms.20-0339

16. Fair J, Paul E, Jones J, Eds. Guidelines to the Use of Wild Birds in Research. Washington, D.C.: Ornithological Council. 2010 August [Cited 2021 August 2] Available from: https://www.ndsu.edu/fileadmin/research/documents/IACUC/Ornithology_Guidelines_August2010.pdf

17. Baker K. Identification guide of European Non-Passerines. Norfolk British: Trust for Ornithology; 2016.

18. Cakmak E, Peksen CA, Bilgin CC. Comparison of three different primer sets for sexing birds. J Vet Diagn. 2017; 29: 59–63. https://doi.org/10.1177/1040638716675197

19. Ushine, N, Kato T, Hayama S. Sex identification in Japanese birds using droppings as a source of DNA. Bulletin of the Japanese Bird Banding Association. 2016; 28: 51–70. (In Japanese) https://doi.org/10.14491/jbba.MS082

20. Burger J, Gochfeld M. Marine birds as sentinels of environmental pollution. Ecohealth. 2004; 1: 263–274. https://doi.org/10.1007/s10393-004-0096-4

21. Fry DM, Maurer JR. Assessment of lead contamination sources exposing California Condors. Species Conservation and Recovery Program Report, 2003-02, California Department of Fish & Game. 2003.April 3 [Cited 2020 July 18] Available from: https://www.researchgate.net/publication/240610549

22. Swaileh KM, Sansur R. Monitoring urban heavy metal pollution using the House Sparrow (*Passer domesticus*). J. Environ Monit. 2006; 8: 209–213. https://doi.org/10.1007/s00244-003-2195-610.1039/B510635D

23. Martinez-Haro M, Green AJ, Mateo R. Effects of lead exposure on oxidative stress biomarkers and plasma biochemistry in waterbirds in the field. Environ. 2011; 111: 530–538. http://dx.doi.org/10.1016/j.envres.2011.02.012

24. Bedrosian B, Craighead D. Blood lead levels of Bald and Golden Eagles sampled during and after hunting seasons in the Greater Yellowstone Ecosystem. In: Watson RT, Pokras M, Hunt G, editors. Ingestion of Spent Lead Ammunition: Implications for Wildlife and Humans. Idaho: The Peregrine Fund; 2009. pp 219–220. https://doi.org/10.4080/ilsa.2009.0209

25. Ministry of the Economy, Trade and Industry. Guidebook for risk assessment of chemicals. (In Japanese) 2007 July [Cited 2020 May 30] Available from: https://www.meti.go.jp/policy/chemical_management/law/prtr/pdf/guidebook_jissen.pdf

26. Krishnan E, Lingala B, Bhalla V. Low-level lead exposure and the prevalence of gout: an observational study. Ann Intern Med. 2012; 157: 233–241. http://dx.doi.org/10.7326/0003-4819-157-4-201208210-00003

27. Harmon BG. Avian heterophils in inflammation and disease resistance. Poult. 1998; 77: 972–977. https://doi.org/10.1093/ps/77.7.972

28. Sharma JM. Overview of the avian immune system. Vet Immunol Immunopathol. 1991; 30: 13–17. https://doi.org/10.1016/0165-2427(91)90004-v

29. Minias P. Evolution of heterophil/lymphocyte ratios in response to ecological and life-history traits: A comparative analysis across the avian tree of life. J Anim Ecol. 2019; 88: 554–565. https://doi.org/10.1111/1365-2656.12941

30. Totzke U, Fenske M, Hüppop O, Raabe H, Schach N. The influence of fasting on blood and plasma composition of herring gulls (*Larus argentatus*). Physiol. 1999; 72: 426–437. https://doi.org/10.1086/316675

31. Franson JC, Pain DJ. Lead in birds. In Beyer WM, Meador JP, editors. Environmental Contaminants in Biota. North Caroline: Taylor & Francis; 2011. pp. 563–594.

32. Burger J, Gochfeld M. Effects of lead on learning in Herring Gulls: An Avian wildlife model for neurobrhaviaral deficits. Neurotoxicology. 2010; 26: 615–624. https://doi.org/10.1016/j.neuro.2005.01.005

33. Fritsh C, Jankowiak L, Wysocki D. Exposure to Pb impairs breeding success and is associated with longerlifespan in under Europran bikachbirds. Sci Rep. 2019; 9: 1–12. https://doi.org/10.1038/s41598-018-36463

34. Haig SM, D’Elia J, Eagles-Smith C, Fair JM, Gervais J, Herring G, Rivers JW, Schulz JH. The persistent problem of lead poisoning in birds from ammunition and fishing tackle. Condor. 2014; 116: 408–428. https://doi.org/10.1650/CONDOR-14-36.1

35. Fenga C, Gangemi S, Di Salvatore V, Falzone L, Libra M. Immunological effects of occupational exposure to lead. Mol Med Rep. 2017; 15: 3355–3360. https://doi.org/10.3892/mmr.2017.6381

36. Cao JJ. Early life exposure to toxic environments: effects on lung and immune cell development in mice and men, University of Groningen, 2016 May 30 [Cited 2020 May 1] Available from: https://www.semanticscholar.org/paper/Early-life-exposure-to-toxic-environments-%3A-effects-Cao-Song/4df8f5642da321ff6ed41554b1f696828cde5f33

37. Arstila TP, Vainio O, Lassila O. Central role of CD4+ T cells in avian immune response. Poult. 1994; 73: 1019–1026. https://doi.org/10.3382/ps.0731019

38. Fang L, Zhao F, Shen X, Ouyang W, Liu X, Xu Y, et al. Pb exposure attenuates hypersensitivity in vivo by increasing regulatory T cells. Toxicol Appl Pharmacol. 2012; 265: 272–278. https://doi.org/10.1016/J.TAAP.2012.10.001

39. Vallverdú-Coll N, Mateo R, Mougeot F, Ortiz-Santaliestra ME. Immunotoxic effects of lead on birds. Sci Total Environ. 2019; 689: 505–515. https://doi.org/10.1016/j.scitotenv.2019.06.251

40. Nain S, Smits JEG. Subchronic lead exposure, immunotoxicologyand increased disease resistance in Japanese quail (*Corturnix coturnix japonica*). Ecotoxicol. 2011; 74: 787–792. https://doi.org/10.1016/j.ecoenv.2010.10.045

41. Centers for Disease Control and Prevention (CDC). Adult blood lead epidemiology and surveillance (ABLES). Cincinnati, OH: US Department of Health and Human Services, CDC, National Institute for Occupational Safety and Health. 2012 February 23 [Cited 2021 December 4] Available from: http://www.cdc.gov/niosh/topics/ables/description.html.

42. Centers for Disease Control and Prevention (CDC). Childhood lead poisoning prevention. Cincinnati, OH: US Department of Health and Human Services, CDC, National Institute for Occupational Safety and Health. 2021 October 27 [Cited 2021 December 3] Available from: https://www.cdc.gov/nceh/lead/data/blood-lead-reference-value.htm

43. Prüter H, Franz M, Auls S, Czirják GA, Greben O, Greenwood AD, Lisitsyna O, Syrota Y, Sitko J, Krone O. Chronic lead intoxication decreases intestinal helminth species richness and infection intensity in mallards (*Anas platyrhynchos*). Sci Total Environ. 2018; 644: 151–160. https://doi.org/10.1016/j.scitotenv.2018.06.297

44. Bichet C, Scheifler R, Cœurdassier M, Julliard R, Sorci G, Loiseau C. Urbanization, Trace Metal Pollution, and Malaria Prevalence in the House Sparrow. PLoS ONE. 2013; 8: e53866. https://doi.org/10.1371/journal.pone.0053866

